# Chronic release of tailless phage particles from *Lactococcus lactis*

**DOI:** 10.1101/2021.07.21.453303

**Authors:** Yue Liu, Svetlana Alexeeva, Herwig Bachmann, Jesús Adrián Guerra Martínez, Nataliya Yeremenko, Tjakko Abee, Eddy J. Smid

## Abstract

*Lactococcus lactis* strains residing in the microbial community of a complex dairy starter culture named “Ur” are hosts to prophages belonging to the family *Siphoviridae*. *L. lactis* strains (TIFN1 to TIFN7) showed detectable spontaneous phage production and release (10^9^-10^10^ phage particles/mL) and up to 10-fold increases upon prophage induction, while in both cases we observed no obvious cell lysis, typically described for the lytic life cycle of *Siphoviridae* phages. Intrigued by this phenomenon, we investigated the host-phage interaction using strain TIFN1 (harboring prophage proPhi1) as a representative. We confirmed that during the massive phage release, all bacterial cells remain viable. Further, by monitoring phage replication *in vivo*, using a green fluorescence protein reporter combined with flow cytometry, we demonstrated that the majority of the bacterial population (over 80%) is actively producing phage particles when induced with mitomycin C. The released tailless phage particles were found to be engulfed in lipid membranes, as evidenced by electron microscopy and lipid staining combined with chemical lipid analysis. Based on the collective observations, we propose a model of phage-host interaction in *L. lactis* TIFN1, where the phage particles are engulfed in membranes upon release, thereby leaving the producing host intact. Moreover, we discuss possible mechanisms of chronic, or non-lytic release of LAB *Siphoviridae* phages and its impact on the bacterial host.

## Introduction

Bacteriophages are highly diverse in shape, structure and composition. They can be icosahedral, spherical, pleomorphic, filamentous, droplet-, bottle- and spindle-shaped; some are with a long or short tail, some are tailless, engulfed in a lipid bilayer or containing lipids beneath the protein capsid; the genetic material can be double-stranded or single-stranded, DNA or RNA (1, 2). The broad accessibility of high-throughput sequencing technologies also revealed a high degree of genetic diversity in bacteriophages; mosaic genomes and numerous novel sequences of unknown function have been reported (3–6). Over 90% of reported phages are tailed double-stranded DNA phages belonging to the order *Caudovirales* (7). Tailed phages primarily interact with their host cell by using tail fibers and baseplate structures, and use the tail for penetrating the bacterial cell surface and viral DNA injection (8, 9). At the end of infection cycle, virulent tailed phage particles are released from the cells by holin-lysin induced lysis of the host. So called temperate bacteriophages undergo an alternative, lysogenic cycle in which the bacteriophage DNA integrates into the chromosome of the host becoming a prophage (10, 11). In this dormant state the prophage can replicate its genome as a part of the bacterial chromosome. Under conditions insulting its host’s DNA integrity the prophage can enter the lytic cycle meaning that it excises from the bacterial chromosome, replicates its genome, assembles into mature phage particles and escapes the host following phage holin-lysin induced cell burst (12, 13).

About 4% of the described bacteriophages lack genes encoding tail proteins and they represent polyhedral, filamentous or pleomorphic phages. Tailless phages use other attachment devises, such as protein complexes or spikes at exposed surface sites (14–16). Some members of this group also apply alternative strategies to release their progeny from infected bacteria. Filamentous phages of the *Inoviridae* family are assembled at the cell surface and excreted from infected cells continuously by extrusion, a process mediated by membrane translocation and channel proteins and that leaves the host cells fully viable (17, 18). Another distinct mechanism of progeny release is budding, a delicate mechanism typical for animal viruses. During budding these viruses are encapsulated by the cell membrane and released, without killing the host. So far, budding has been suggested only for the family of *Plasmaviridae*, tailless phages infecting wall-less bacteria *Acholeplasma* species via membrane fusion (19, 20). In contrast to lytic phage release that kills the host, the non-lytic release is also referred to as chronic release (21). The group of tailless phages includes bacteriophages that have, in addition to nucleic acid and proteins, internal or external lipid constituents - a property originally associated with viruses infecting multicellular eukaryotes. Currently, the lipid-containing bacteriophages are classified into four families, *Corticoviridae*, *Cystoviridae*, *Plasmaviridae* and *Tectiviridae* (22).

Notably, all currently known phages infecting lactic acid bacteria (LAB) are members of the *Caudovirales* order (7), or tailed phages. Bacterium-phage interactions play a key role in the evolution of both partners in the interaction. Earlier, we described (pro)phages abundantly released and co-existing within a naturally evolved microbial community – mixed (originally undefined) complex starter culture of LAB used in dairy fermentations (23). These cultures represent an interesting model ecosystem because it was established through long term propagation by back-slopping. Practicing back-slopping creates the boundaries for natural selection which drives adaptive evolution of the culture and its constituent microbial strains.

Based on analysis of the genomic content the isolated (pro)phages belong to P335 group lactococcal phages of *Siphoviridae* family, order *Caudovirales*. However, they possess some peculiar features: phage particles are abundantly released spontaneously and further stimulated by mitomycin C induction (23). They appear to be tailless due to disruptions in tail-protein encoding genes (6). Moreover, the release of the (pro)phages from the host cells was not accompanied by detectable cell lysis, a phenomenon which is typical for release of *Siphoviridae* bacteriophages (24–27). We set out to investigate this phenomenon in this study. Here we demonstrate that the tailless *Siphoviridae* phage particles are enclosed in lipid membrane and are released from the cells by a non-lytic mechanism, a phenomenon not described before in LAB phages.

## Materials and methods

### Strains and media

All *Lactococcus lactis* strains used in this study were statically cultivated in M17 broth (OXOID) with 0.5% (wt/vol) lactose addition (OXOID) at 30°C, unless specified otherwise. All *Escherichia coli* strains harboring plasmids used in this study were cultivated at 37°C, in LB broth (BD Difco) supplemented with 150μl/ml erythromycin and shaken at 120 rpm.

### Cell growth, prophage induction, phage purification and quantification

Overnight cultures in M17 broth supplemented with 0.5% lactose (LM17) were diluted up to OD_600 nm_ = 0.2 and allowed to grow for 1 hour at 30ºC before mitomycin C (MitC) was added (final concentrations of 1 μg/ml). For control purposes, the same diluted cultures without MitC were used. Incubation proceeded for 6 or 7 hours and the turbidity at 600 nm was monitored at 1 hour intervals. At the end of induction the total cell number was determined by direct counting in a haemocytometer chamber and the viable count was made by a standard spread plating on M17 agar supplemented with 0.5% lactose. Released phage particles were concentrated from the culture supernatants by PEG/NaCl precipitation as previously described (6), and the quantity was estimated based on phage DNA content in culture supernatants or in PEG/NaCl concentrated phage suspensions using agarose gel electrophoresis as previously described (23).

### Construction of plasmids for chromosomal integration into prophage sites

Plasmid pSA114 is a derivative of plasmid pCS1966 (28) - the chromosomal integration vector, allowing positive selection of cells in which the plasmid had been excised from the genome, resulting in unmarked integrations in the chromosome of *L. lactis*.

Two DNA fragments 671 and 941bp of adjacent loci of prophage (proPhi1) were amplified from *L. lactis* TIFN1 chromosome using 1M_HR1_Fw+/1M_HR1_Rv and 1M_HR2_Fw/1M_HR2_1Rv+ primer pairs respectively (see table 1). The two fragments were interconnected by multiple cloning site (MCS) introduced by PCR overlap extension mutagenesis in order to allow further insertions between the homology arms. The resulting 1.7 Kb-PCR fragment was digested with KpnI and NcoI and ligated into corresponding sites of pSEUDO-GFP (29) resulting in plasmid pSA114. The 34 bp MCS between the amplified prophage sequences of pSA114 was used for further cloning. pSA116, the vector for integration of CmR (chloramphenicol resistance cassette, *cat*) into prophage was made by inserting CmR between the prophage homology regions of pSA114. Chloramphenicol cassette (*cat*) was amplified by PCR using pGhCAM2_Fw/pGhCAM_Rv primers (Table 1) and pVE6007 [pGhost7, (30)] as a template. The fragment was digested with EcoRI and BamHI and sub cloned into corresponding sites of pSA114, yielding pSA116. Plasmid pSA120, the vector for integration of *gfp* [the gene of the superfolder variant of GFP, (31)] into prophage was made as follows. The *gfp* flanked by CP25 artificial promoter (32) and two terminator sequences was excised from pIL-JK2 using EcoRI and BamHI and inserted into the same site (between the prophage homology regions) of pSA114, yielding pSA120.

**Table 1.**
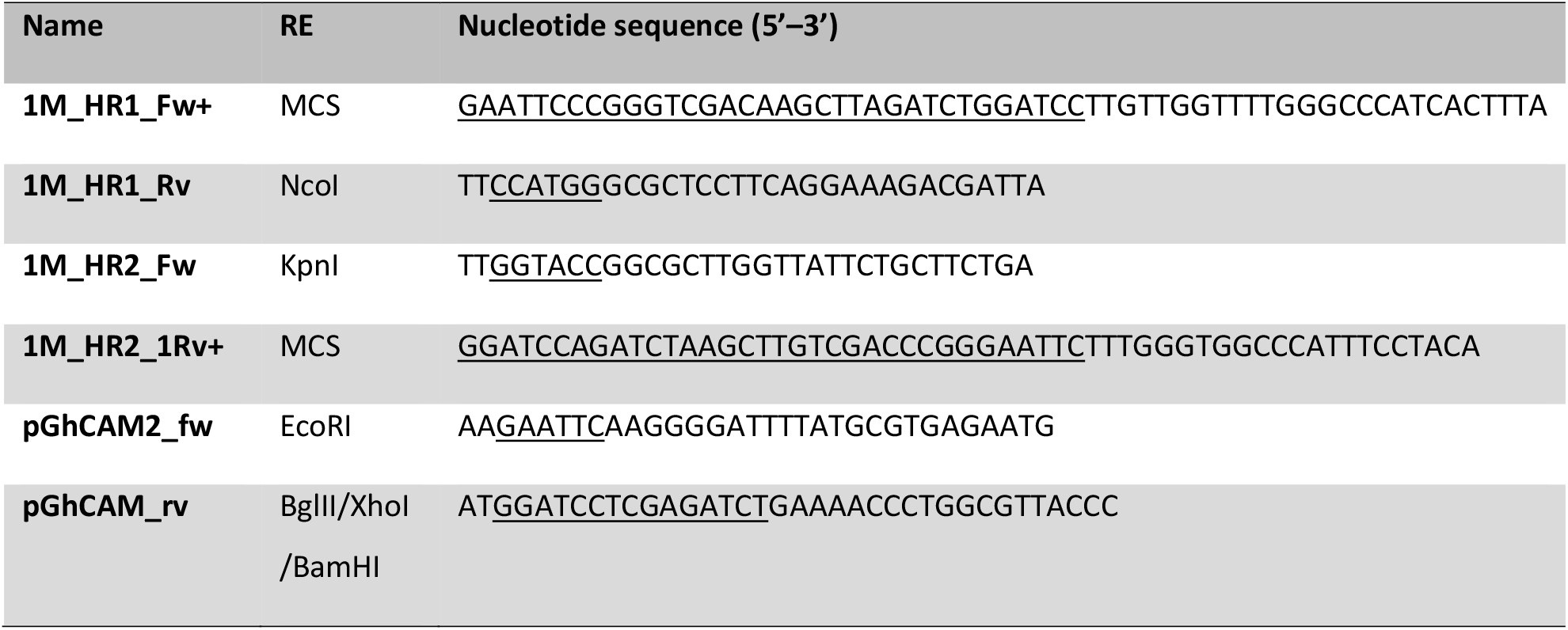
Primers used in this study. Restriction enzyme (RE) sites are underlined.

### Modification of the chromosomal integration strategy for industrial strains and construction of new integration vectors

The designed vectors are derivatives of pCS1966 and unable to replicate in *L. lactis*. The original strategy (28) includes both transformation and chromosomal integration by homologous recombination for the successful acquisition of such vectors by *L. lactis* cells. Therefore, the plasmid acquisition is drastically dependent on the efficiency of transformation. Industrial “wild” strains are usually featured by poor transformability compared to “domesticated” laboratory *L. lactis* strains. Therefore, a new strategy has been designed by splitting plasmid transformation and its homologous recombination events. We constructed vectors that combine *L. lactis* thermosensitive (Ts) replication origin *repA^Ts^* and *oroP* gene. Ts replication origin allows the plasmid replication under permissive temperature after the transformation event, followed by integration through homologous recombination at elevated temperature. Gene *oroP* enables counterselection for loss of the plasmid backbone, leaving unmarked integrations in the chromosome of *L. lactis* at specific target sites.

New vectors, pSA130-YL and pSA132-YL were constructed as follows. *E. coli* strain EC1000 (33) was used for cloning and plasmid propagation. pG^+^host9 (30) was used to provide the backbone with *repA^Ts^* and *ermAM*. The plasmids pSA116 and pSA120 were used to provide the cassettes of DNA labels (CmR or *sf-gfp*) flanked by proPhi1 homology regions (MHR) and *oroP*. The KpnI/FspI fragment of pSA116, was ligated into KpnI/EcoRV digested pG^+^host9, resulting in pSA130-YL. pSA132-YL was constructed in two steps: first KpnI/SalI fragment of pSA120 was ligated into the corresponding site of pG^+^host9, then SalI/FspI fragment of pSA120 was ligated into SalI/EcoRV site, yielding pSA132-YL.

### Plasmid integration and backbone elimination

*L. lactis* transformation was performed using a modified protocol as we described earlier (23). The confirmed transformants were propagated at the permissive temperature, 28°C in M17 broth (0.5% glucose or lactose) with 3 μg/ml erythromycin, and stored in 15% glycerol at −80°C until further use. For the integration step, *L. lactis* cells transformed with constructed plasmids were incubated at 37°C overnight, the OD_600_ of cultures was measured, cells were plated in proper dilutions (depending on OD_600_ values) on selection plates (L/GM17, 1.5% agar, 0.5 M sucrose and 3 μg/ml erythromycin), and incubated at 37°C till colonies emerged. The presence and orientation of the whole plasmid inserts were checked with PCR, and correct validated clones were maintained as 15% glycerol stocks at −80°C.

For the backbone elimination step, the validated clones with integrated plasmids were inoculated in 2 ml SA medium (34) containing 1% lactose or glucose and incubated at 30°C overnight. Then they were diluted 10x in SA (1% lactose or glucose) medium and incubated 30°C for 6 h. Thereafter 10 μl of the culture was streaked on SA (1% lactose or glucose) agar plate supplemented with 10 μg/ml 5-fluoroorotate (Sigma). Plates were incubated at 30°C until 5-fluoroorotate resistant colonies emerged. For resulting labelled strains, TIFN1::*cat* and TIFN1::*gfp* (inserted genes from corresponding plasmid-donors, pSA130-YL and pSA132-YL), correct inserts and their location on TIFN1 chromosome were confirmed by PCR, sequencing the PCR products and phenotypic analysis: either green fluorescence or chloramphenicol resistance.

### Phage lipid and DNA labelling

The dyes used for labelling of lipids and DNA are shown in Table 2. To stain DNA, PEG precipitated phage particles were incubated with Sybr Green (Invitrogen, Molecular probes Cat. no. S7563) at 80°C for 10 minutes in 10^−4^ final dilution of commercial stock as described earlier (35) or with GelRed nucleic acid gel stain (Biotium,10^−4^ final dilution of commercial stock) under the same conditions.

**Table 2.** Corresponding labelling of lipid and DNA dyes.

| Label used in text | Name of the dye |
| --- | --- |
| **Lipophilic dye 1** | FM 4-64 |
| **Lipophilic dye 2** | MitoTracker Green FM |
| **Lipophilic dye 3** | CellMask Deep Red |
| **DNA dye 1** | Sybr Green |
| **DNA dye 2** | GelRed Nucleic Acid Gel stain |

For membrane detection CellMask DeepRed (Life Technologies GmbH) was used in final dilution 10^−3^ of commercial stock and phages were incubated for 5 minutes at 37°C; FM4-64 (Molecular Probes) was used in a final concentration 20 μg/ml, incubation proceeded 15 minutes at room temperature; and MitoTracker Green FM (Molecular Probes Cat. no. M7514) was used at final concentration of 20 nM and phages were incubated with the dye for 15 minutes at 37°C. For double staining of DNA and membrane, MitoTracker was added to the phages stained with GelRed, the samples were vortexed and measured immediately by flow cytometry. In control experiment the membranes were first extracted by adding to the phage suspension equal volumes of chloroform, the samples were vortexed, centrifuged during 3 minutes at 14000× g, aqueous phase containing the phage particles was collected and the staining was performed as described above.

When phages were not added, no detectable fluorescent particles were present in the control samples. To exclude contamination of phage suspension by bacterial cells, bacteria were added to phage suspension prior to staining of either DNA or lipids. In these control samples an additional population of particles with much higher fluorescence intensity was detected (not shown), as anticipated, given a bacterial cell contains much higher amounts of lipids and DNA per particle compared to phages.

### Flow Cytometry

Prior to flow cytometry analysis 2 μl of fluorescent microspheres (1×10^−3^ of the stock Fluoresbrite^®^ YG Microspheres 0.75μm, Polysciences) was added and the volume was adjusted to 500 μl by adding FACSFlow solution (10 mM phosphate-buffered saline, 150 mM NaCl, pH 7.4; Becton-Dickinson).

Samples were analyzed by using a BD FACS Aria™ III flow cytometer (BD Biosciences, San Jose, CA). The cytometer was set up using an 85 μm nozzle and was calibrated daily using BD FACSDiva Cytometer Setup and Tracking (CS&T) software and CS&T Beads (BD Biosciences). An 488 nm, air-cooled argonion laser and the photomultipliers with 488/10 band pass filter for forward and side scatter and with filter 530/30 nm (with 502 LP filter) was used for the detection of GFP, Sybr Green and MitoTracker. GelRed was excited with a yellow-green 561nm laser and detected using a 610/20 nm with LP 600 nm filter. CellMask DeepRed was excited with 633 nm laser and detected with 660/20 nm filter. FM4-64 dye was excited with 561 nm laser and detected with a 780/60 nm with LP 735 nm filter. FSC and SSC voltages of 300 and 350, respectively, and a threshold of 1,200 on FSC was applied to gate on the bacteriophages and bacterial cells population.

Data were acquired by using BD FACSDiva™ software and analyzed by using FlowJo flow cytometry analysis software (Tree Star, Ashland, OR).

### Chemical lipid analysis

Normal phase high performance liquid chromatography (NP-HPLC) with evaporative light scattering detection (ELSD) was used for the quantitative analysis of phospholipids (36).

The analyses were performed with a 600E System Controller (Waters), vacuum degasser (Knauer), 231 XL sampling injector (Gilson) and a 3300 (ELSD) evaporative light scattering detector (Alltech). Extraction of the phospholipids from 2 g freeze dried milk sample was done with a mixture of chloroform, methanol and ammonia (NH_3_) in water. After centrifugation of the sample 10 min at 4500 g, 10.0 mL of the supernatant was evaporated under vacuum at 40°C in a heating block. When the sample was dried, 1 mL absolute ethanol was added and again evaporated to dryness. The dried sample was dissolved in 1.0 mL of the phospholipid solvent containing iso-octane, chloroform and methanol.

Fifty milliliter of the sample solution was injected on an Xbridge amide analytical column, 3.5 μm, 4.6 × 250 mm (Waters). The components were eluted at a flow rate of 1.0 mL/min with a gradient of eluent A (iso-octane and acetone) and eluent B (2-propanol and ethyl acetate) to eluent C (2-propanol, water, ammonia and acetic acid) in 50 minutes. Data analysis was done with Chromeleon software version 7.2 (Thermo Fisher Scientific). The non-linear response of the ELSD was converted to a more linear signal in order to increase the accuracy of the quantification of phospholipids differing in fatty acid composition compared to those of the standard.

1,2-dipalmitoyl-sn-glycero-3-phosphoryl ethanolamine (PE, Matreya), 1,2-dipalmitoyl-sn-glycero-3-phosphoryl glycerol (PG, Matreya), 3-sn-lyso phosphatidyl ethanolamine (LPE, Sigma), DL-α-phosphatidyl choline (dipalmitoyl, C16:0) (PC, Sigma), Sphingomyelin (SM, Sigma), phosphatidyl serine (oleoyl) (PS, Matreya), lyso-phosphatidylcholine (palmitoyl) (LPC, Matreya) and phosphatydyl inositol (linoleoyl) (PI, Matreya) were used as calibration standards for quantitative analysis. A reference sample (buttermilk powder) with known amounts of phospholipids was analyzed and recovery of spiked phospholipids was performed to control for accuracy and precision of the method.

### Scanning/transmission electron microscopy

For scanning electron microscopy, *L. lactis* TIFN1 and TI1c cultures were subjected to 1 μg/ml MitC induction as described above. After 5h of induction the samples were fixed with 2.5% glutaraldehyde in PBS buffer for 1 hour at room temperature. A droplet of the fixed cell suspension was placed onto poly-L-lysine coated coverslips (Corning BioCoat, USA) and allowed to stand for 1 hour at room temperature. After rinsing in PBS the samples were post stained in 1% osmium tetroxide in PBS. Subsequently the samples were dehydrated in a graded series of ethanol followed by critical point drying with CO_2_ (Leica EM CPD300, Leica Microsystems). The coverslips were fitted onto sample stubs using carbon adhesive tabs and sputter coated with 6 nm Iridium (Leica SCD500). Samples were imaged at 2 KV, 6 pA, at room temperature in a field emission scanning electron microscope (Magellan 400, FEI Company, Oregon, USA).

For transmission electron microscopy, purified phage particles were subjected to negative staining and examined exactly as descried previously (23).

## Results

### No detectable cell lysis during phage release

A previous study on *Lactococcus lactis* strains TIFN1-7 originating from the mixed cheese starter culture indicated no obvious drop in the optical density of the bacterial cultures following prophage activation (23). We were triggered by this observation and therefore we further examined this phenomenon using strain TIFN1 as a model. We first analyzed the cell viability in prophage induced cultures supplemented with mitomycin C (MitC) and control cultures without MitC induction. The total cell counts determined using a haemocytometer and the number of culturable cells, i.e., colony forming units, were similar (supplementary Fig. S1), indicating that at least the vast majority of TIFN1 cells present in the tested conditions remained viable. Both cell counting methods showed no obvious differences in cell numbers from the prophage-induced cultures and non-induced control cultures, further confirming that in the *L. lactis* strains, represented by lysogen TIFN1, there was no detectable cell lysis in spite of the abundant phage release upon phage induction.

### The major part of the culture actively produces phages

To elucidate whether phage production is a population-wide activity in a clonal culture of strain TIFN1, we monitored *in vivo* phage replication using a reporter strain, in which a *gfp* reporter was inserted within the prophage. In cells actively replicating the phage particles, green fluorescence intensity was expected to increase. As mentioned, we used *L. lactis* TIFN1 as the model strain, which harbors the genome of prophage proPhi1 (6). The insertion site was selected within the prophage sequence between stop codons of open reading frames (ORFs) 48 and 49 encoded on opposite DNA strands (Fig. 1A), resulting in strain TIFN1::*gfp*. In parallel, a fluorescence-negative control strain was constructed in which the chloramphenicol resistance gene *cat* was inserted at the same site, yielding strain TIFN1::*cat*.

**Figure 1.**
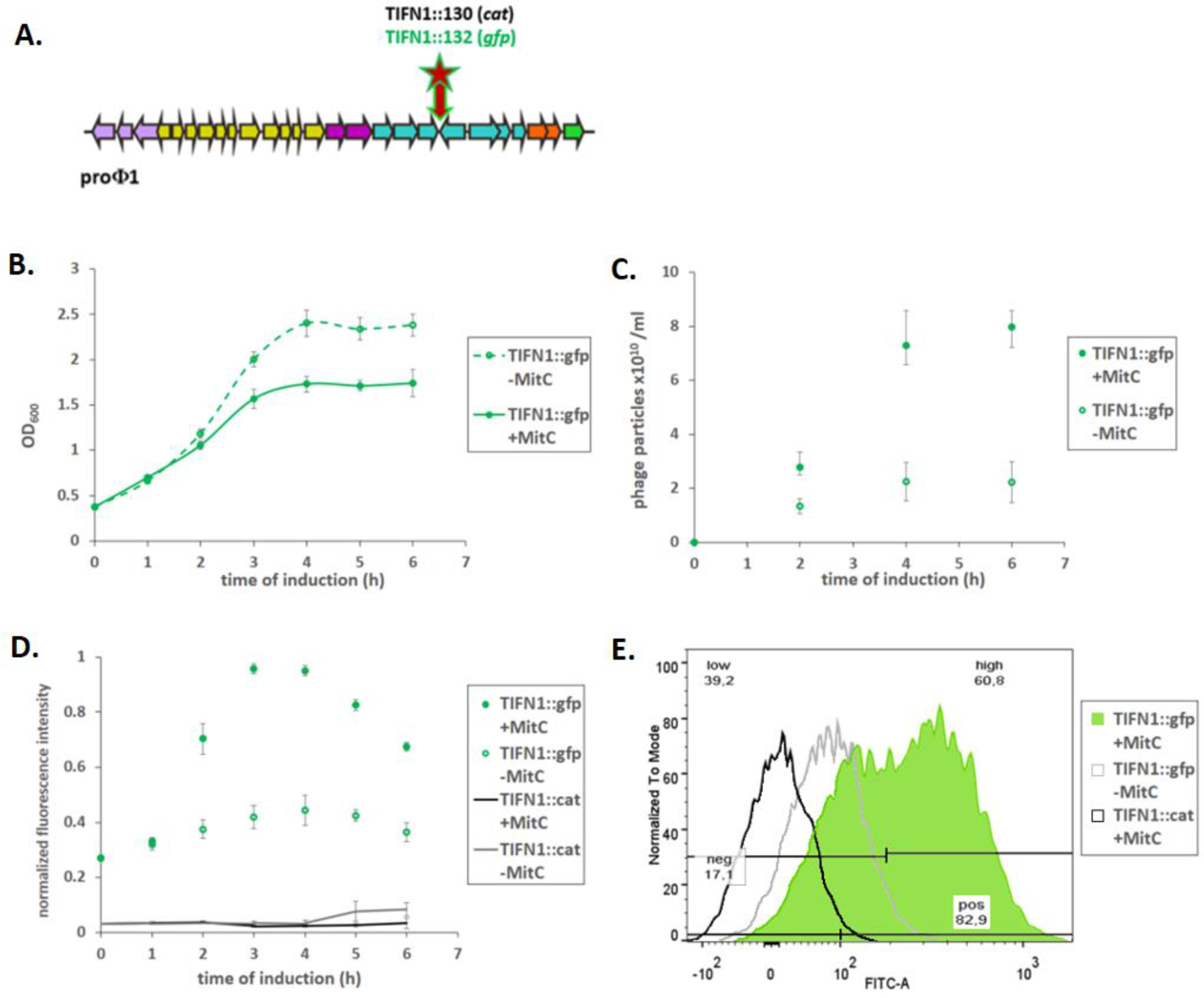
Phage labelling and examination of phage replication. **(A)** Schematic drawing of the prophage genome with marked *gfp* and *cat* insertion sites. Arrows represent ORFs and indicate the direction of gene transcription. The number of arrows does not reflect the real ORF numbers but is only a schematic presentation. The insertion was made in between two convergent ORFs. Colors in arrows schematically represent different phage gene clusters. **(B)** Growth response of TIFN1::*gfp* to MitC treatment. **(C)** Phage release by TIFN1::*gfp* during MitC induction. Green symbols represent MitC treated cultures, grey symbols represent control cultures without MitC. **(D)** Dynamics of phage replication (as derived from average cell fluorescence intensity) during MitC induction (green symbols) and in uninduced samples (grey symbols) in reporter strain (TIFN1::*gfp*) compared to base-line fluorescence of non-*gfp* cultures (TIFN1::*cat*, black and grey lines for induced and uninduced conditions respectively). **(E)** Fluorescence distribution in the population at 3 hours of induction in non-*gfp* TIFN1::*cat* (black unfilled), uninduced TIFN1::*gfp* (grey unfilled), and MitC induced TIFN1::*gfp* (green filled) cultures. The statistics in (e) is shown for induced TIFN1::*gfp*: 82.9% of the population was positive for green fluorescence (pos 82.9), 17.1% was fluorescence-negative (neg 17.1); 60.8% was highly fluorescent (high 60.8) and 39.2% was low in fluorescence (low 39.2).

The derived strain TIFN1::*gfp* showed similar growth behavior (Fig. 1B) to the wild-type TIFN1 (data see Alexeeva et al., 2018) with and without prophage induction by MitC, as indicated by monitoring culture turbidity. TIFN1::*gfp* also produced phage particles (Fig. 1C) to a similar amount as the wild-type (data see Alexeeva et al., 2018): 10^10^ phage particles/mL were found in cultures without added MitC and phage numbers increased to ~10^11^/ml upon MitC induction in 6 h, as estimated by quantifying phage DNA content.

To study the *in vivo* dynamics of prophage induction in *L. lactis* TIFN1 we used strain TIFN1::*gfp* and followed in time the fluorescence intensity of the cells by flow cytometry in MitC-induced and un-induced cultures. As a fluorescence-negative control we used TIFN1::*cat*. TIFN1::*cat* exhibited very low background fluorescence, not changing in time and not affected by MitC addition (Fig. 1D). The un-induced culture of TIFN1::*gfp* showed moderate fluorescence already at the initiation of induction (time point 0 h), and the fluorescence increased slightly in time. This is in line with the observed constitutive phage induction and replication taking place even without MitC induction (Fig. 1C and Fig. 1D). The induced culture of TIFN1::*gfp* showed a clear increase in fluorescence intensity till the 4^th^ hour post-induction, and then the fluorescence intensity declined slightly. The increase in fluorescence intensity of the cells was 2.5-3 fold and correlated with the increase in the number of released phage particles (Fig. 1C and Fig. 1D).

To examine whether the major fraction of the bacterial population actively produces phage particles, the distribution of fluorescence in individual cells was measured by flow cytometry. The fluorescence distribution per particle in the negative control (TIFN1::*cat*, black unfilled) as well as in un-induced cultures of TIFN1::*gfp* (grey unfilled) and MitC induced TIFN1::*gfp* (green filled) at time point 3 hours post-induction was measured (Fig. 1E). In the MitC induced TIFN1::*gfp* culture more than 80% of the cells are green fluorescent and more than 60% are highly fluorescent, which is a distinct population (second green peak in Fig. 1E). This indicates that the majority of the cells in the population actively replicate phage DNA and produce phage proteins. When relating this observation to the cell count results (supplementary Fig. S1), where no obvious reduction in cell number was detected under phage inducing conditions, the hypothesis of non-lytic phage release is supported.

### Phages are enclosed in lipid bilayers

Non-lytic, chronic phage release has been previously described to occur via budding (*Plasmaviridae*) or extrusion (*Inoviridae*) (17–20). In case the budding mechanism of cell exit is recruited by the phage particles, it is expected to be enveloped by cellular lipids upon release. Therefore, first of all, we analyzed the presence of a lipid bilayer in/engulfing the phage particles. We employed three lipophilic dyes staining cellular membranes/lipid bilayers, but all essentially non-fluorescent in aqueous media. All three lipophilic dyes were efficiently staining the phage particles (Fig. 2A–C), confirming presence of lipid membranes. Moreover, when the phage particles were treated with chloroform prior to staining with lipophilic dye 3, the fluorescence was largely abolished (Fig. 2C, blue line). The chloroform treated particles were visualized by EM and showed typical morphology of phage heads (see Fig. 1A & B from Alexeeva et al., 2018). We further confirmed that the lipid enclosed particles are indeed bacteriophages containing DNA: the phage particles were readily stained with the two DNA dyes (Fig. 2D & E). Moreover, double staining with DNA dye 2 (red fluorescence) in combination with lipophilic dye 2 (membrane stain, green fluorescence) resulted in double stained particles, confirming that the phage particles indeed contain DNA and are enclosed by membranes (Fig. 2F). This conclusion is also supported by the previous study where the tailless phage particles were isolated with the same method and subjected to DNA sequencing, and full phage genomes were recovered with more than 100-fold higher coverage than background (6).

**Figure 2.**
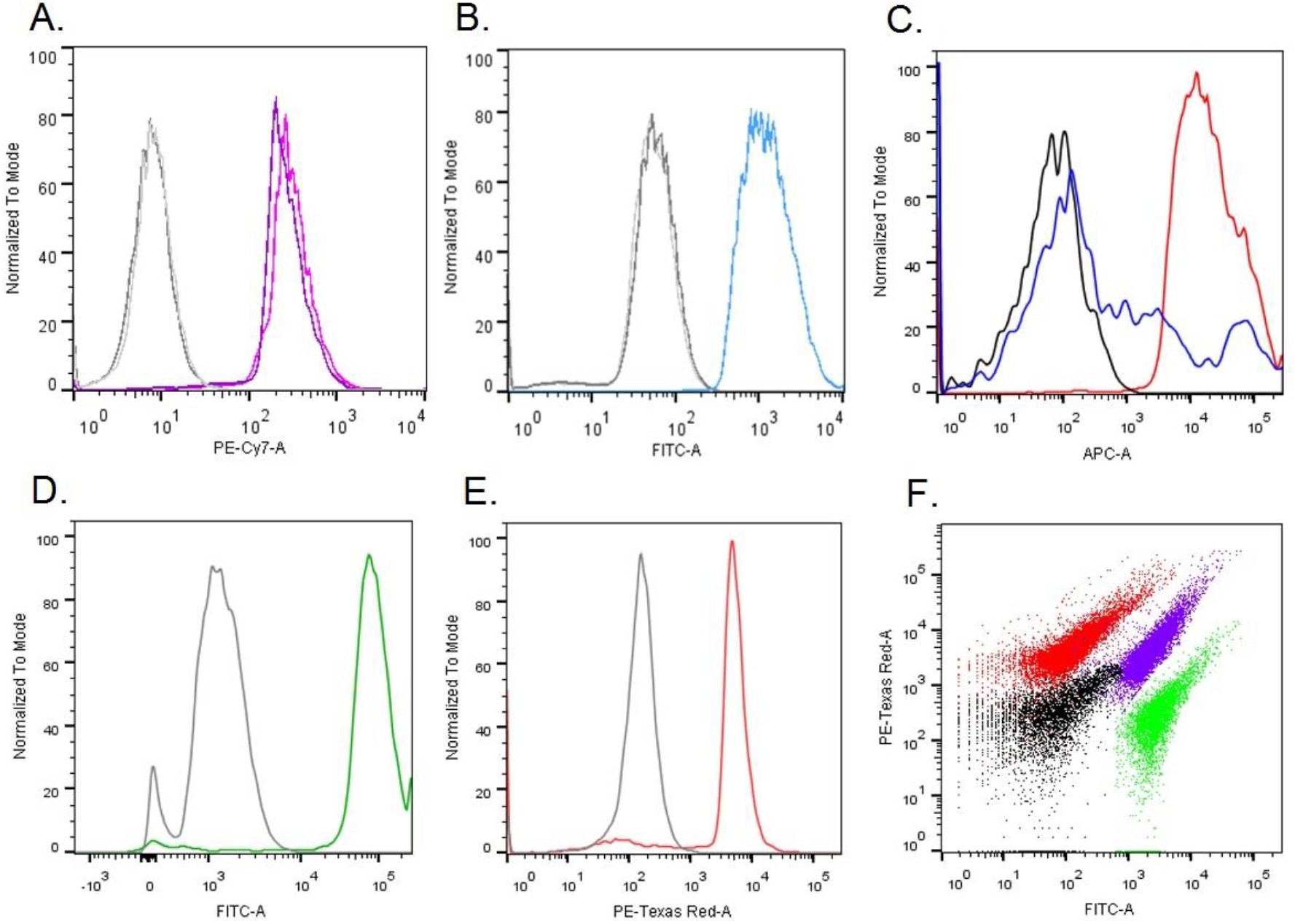
Staining proPhi1 particles with various lipophilic (A, B, C, F) and DNA binding (D, E, F) dyes followed by flow cytometry analysis. Grey/black - unstained phage particles. **(A)** Lipophilic dye 1; **(B)** Lipophilic dye 2; **(C)** Lipophilic dye 3, the blue line represents the sample stained after chloroform treatment; **(D)** DNA dye 1; **(E)** DNA dye 2; **(F)** superimposed dot plot of proPhi1 particle samples with different staining: unstained (black), lipophilic dye 2 (green), DNA dye 2 (red), and double stained lipophilic dye 2 and DNA dye 2 (purple-blue).

Since the hypothesis of phage particles being enclosed in lipid bilayer is now supported by experimental evidence, we continued to find additional support by studying the phage particles with transmission electron microscopy (TEM). In this case, phage particles were not pre-treated with chloroform to retain the lipid membrane, and we compared the particle morphology and size to chloroform-treated phage particles. It was observed that the morphology of untreated (Fig. 3A) and chloroform-treated (Fig. 3B) particles was similar, although they did show different electron-densities as reflected by the different darkness of particles, possibly indicating differences in compositions as chloroform will disintegrate lipid bilayers and dissolve lipids. We also noticed a difference in particle sizes caused by chloroform treatment. When measuring the particle diameters (defined as the distance between two opposite corners of the hexagon shape, measured by ObjectJ), untreated particles showed diameters of 65.4 ± 4.1 nm (n=54), significantly (p<0.00001, 2-tailed, unequal variance) larger than chloroform-treated particles with diameters 58.0 ± 2.0 nm (n=27). The difference in average diameters, 7.4 ± 4.6 nm, coincides with the thickness of two lipid bilayers (37). This analysis also supports the hypothesis that the released phage particles of *L. lactis* TIFN1 are enclosed in lipid bilayers.

**Figure 3.**
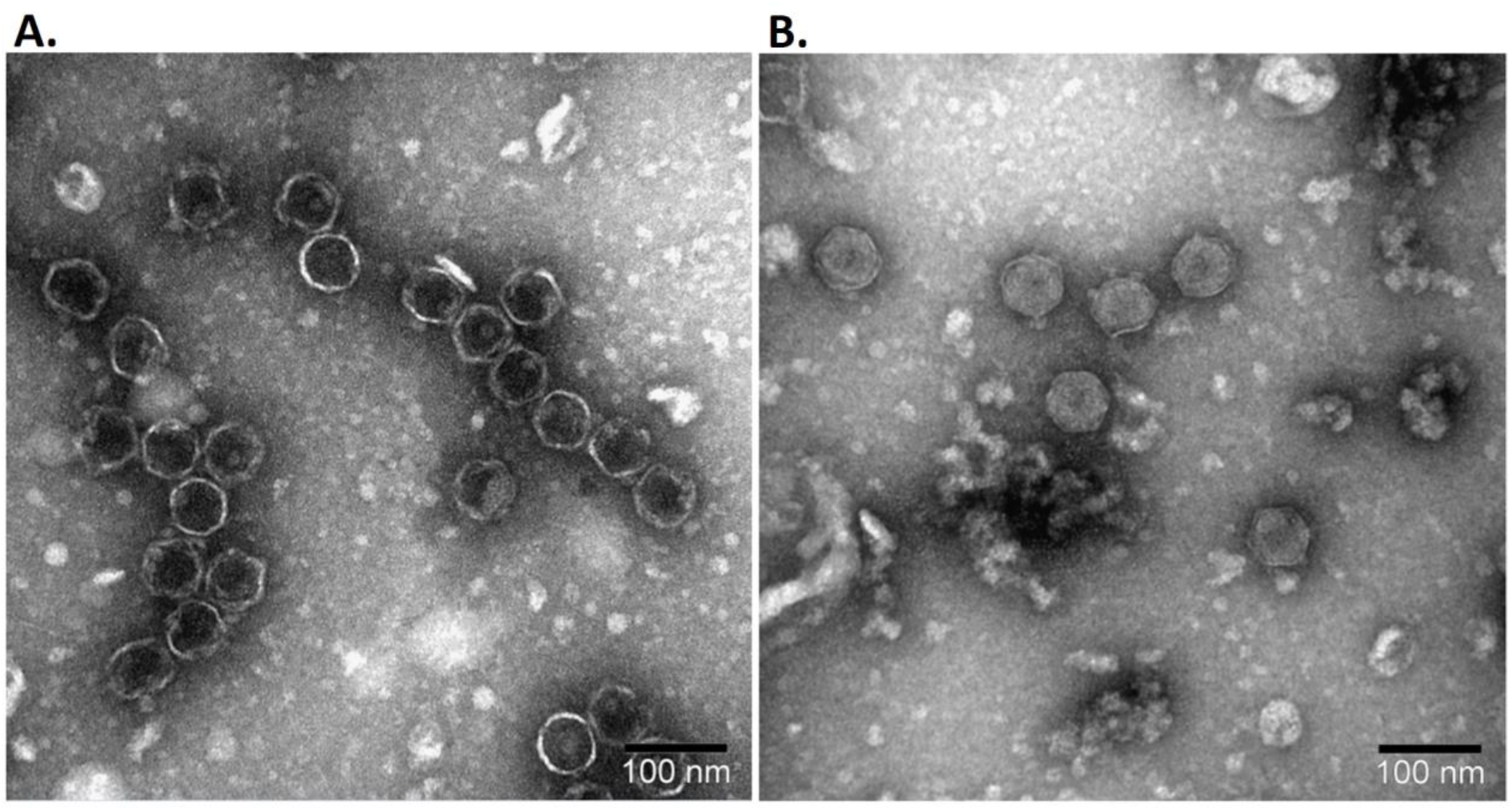
Transmission electron micrograph of proPhi1 with (A) and without (B) chloroform treatment.

### Lipid composition of phage particles differs from host cells

Next, we extracted the lipids from phage crops produced by strain TIFN1 and also from whole cell-derived protoplasts and subjected them to chemical lipid analysis using liquid chromatography coupled with mass spectrometry (LS-MS). As a phage-free control, phage-cured strain TI1c (23) was subjected to the same procedure of prophage induction and purification from the culture supernatant. The TIFN1 phage specimen lipid signals were well above the background level of the phage-free control from TI1c (supplementary Fig. S2). Phosphatidyl glycerol (PG) and cardiolipin (CA) were detected in phage samples as well as in cellular lipid samples, however, the ratio between the two major lipid species differed between the phage and the cell membrane lipid samples (Fig. 4).

**Figure 4.**
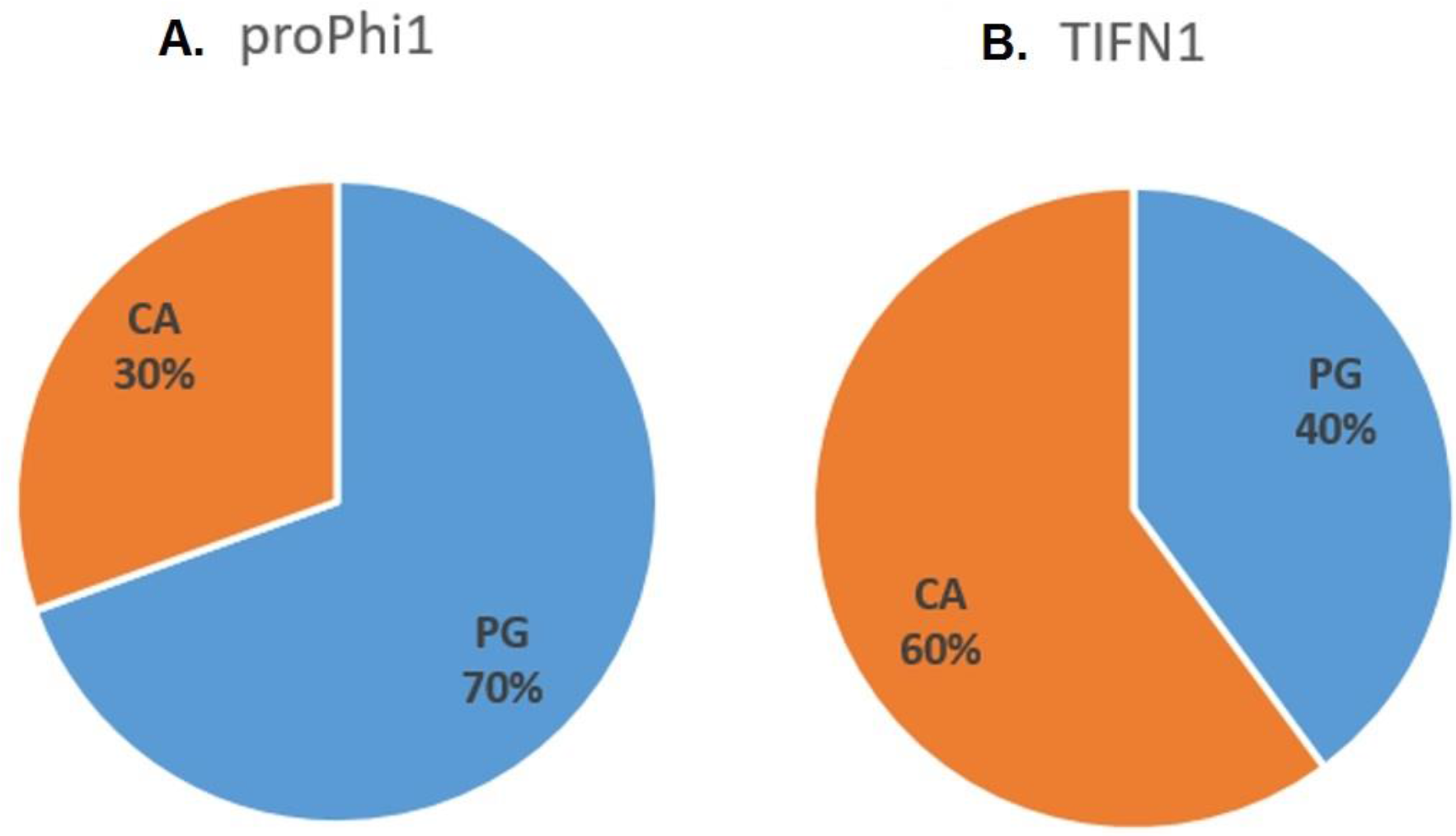
Phage and cell lipid composition. Composition of lipids extracted from A) isolated proPhi1 phage particles and **(B)** TIFN1 whole cell-derived protoplast. PG, phosphatidyl glycerol; CA, cardiolipin.

The major lipid in the *L. lactis* cell membrane is cardiolipin and a CA/PG ratio of about 2.2 has been determined for *L. lactis* membrane earlier (38). We found the CA/PG ratio value of 1.5 for cellular lipids extracted from TIFN1 (Fig. 4B). Remarkably, lipids of the phage crops were enriched in phosphatidyl glycerol with a CA/PG ratio of 0.4 (Fig. 4A). This suggests that the released phage particles are possibly enclosed by phospholipids derived from distinct regions of lipid rafts/domains in the *L. lactis* cell membrane (39, 40).

To further characterize the phage release from the cells we employed scanning electron microscopy to observe MitC induced cells of wild-type strain TIFN1 and its prophage cured derivative strain TI1c (Fig. 5). The MitC treated TI1c had the usual morphology and smooth surface of a Gram-positive coccus without any detectable alteration (Fig. 5C & D). Strain TIFN1 however, showed a ruffled cell surface and accumulated numerous budlike, small spherical structures, typically near the cell division septum (Fig. 5A & B). The phage cured strain TI1c lacked these extracellular structures.

**Figure 5.**
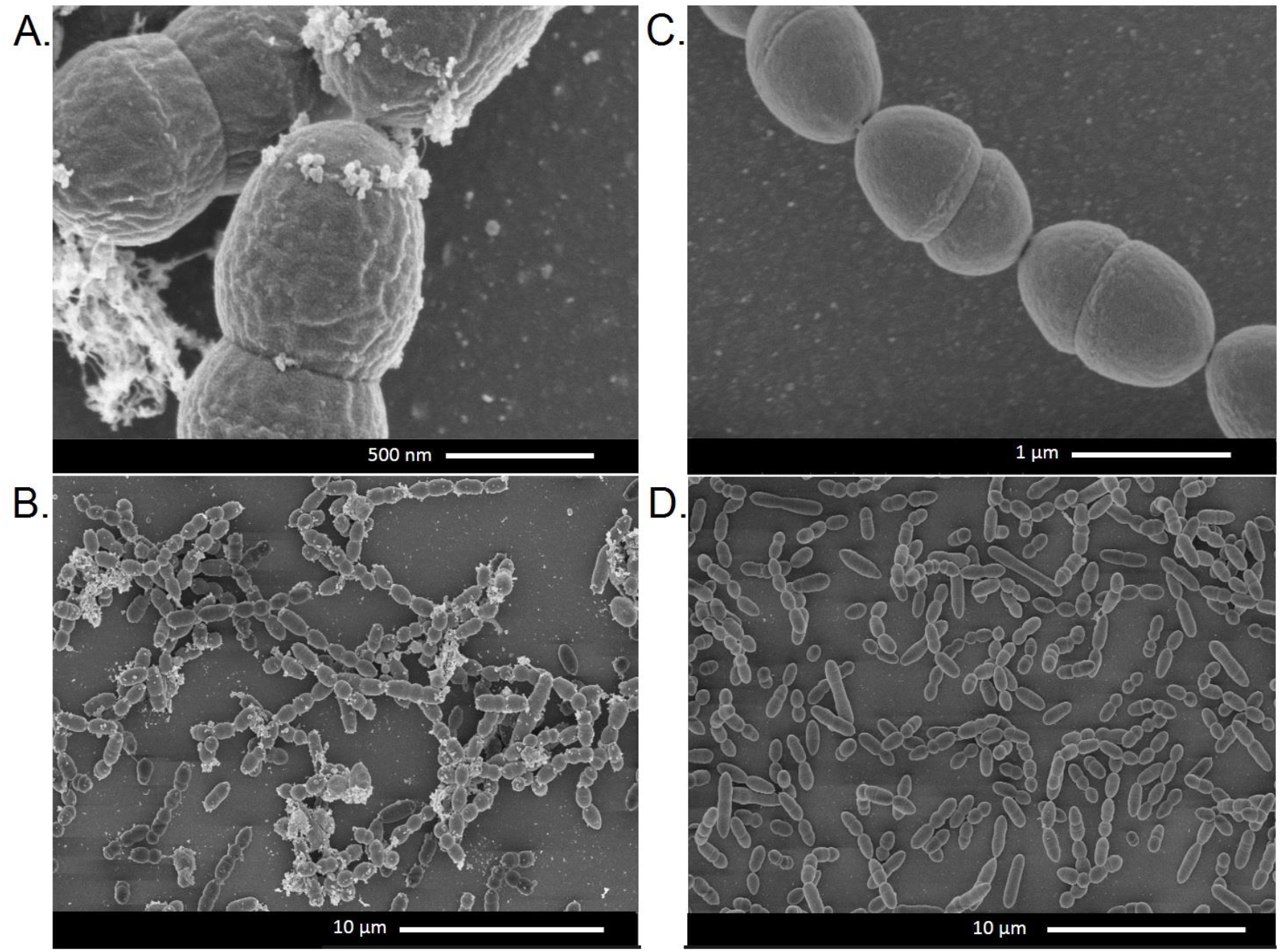
Scanning electron micrograph of cells subjected to 6h MitC treatment. **(A)** and **(B)** TIFN1, **(C)** and **(D)** TI1c.

The observation that extracellular particles are accumulated near the cell division planes is in line with our speculation made from the observation of a difference in lipid composition between phage particles and the host cells indicating that the processes of phage engulfing and release are specific for defined regions of the cell membrane.

## Discussion

Bacteriophages are thought to be the most abundant biological entities on Earth and adopted a striking variety of forms and mechanisms of interaction with their host cells (41). Combining observations from this study, we propose a novel mechanism of interaction for lactococcal phages and their hosts, where the tailless *Siphoviridae* phage particles are enclosed in a lipid membrane and are released from the cells by a non-lytic mechanism (Fig. 6). This chronic, non-lytic phage release mechanism has not been previously described for LAB phages or *Siphoviridae* phages.

**Figure 6.**
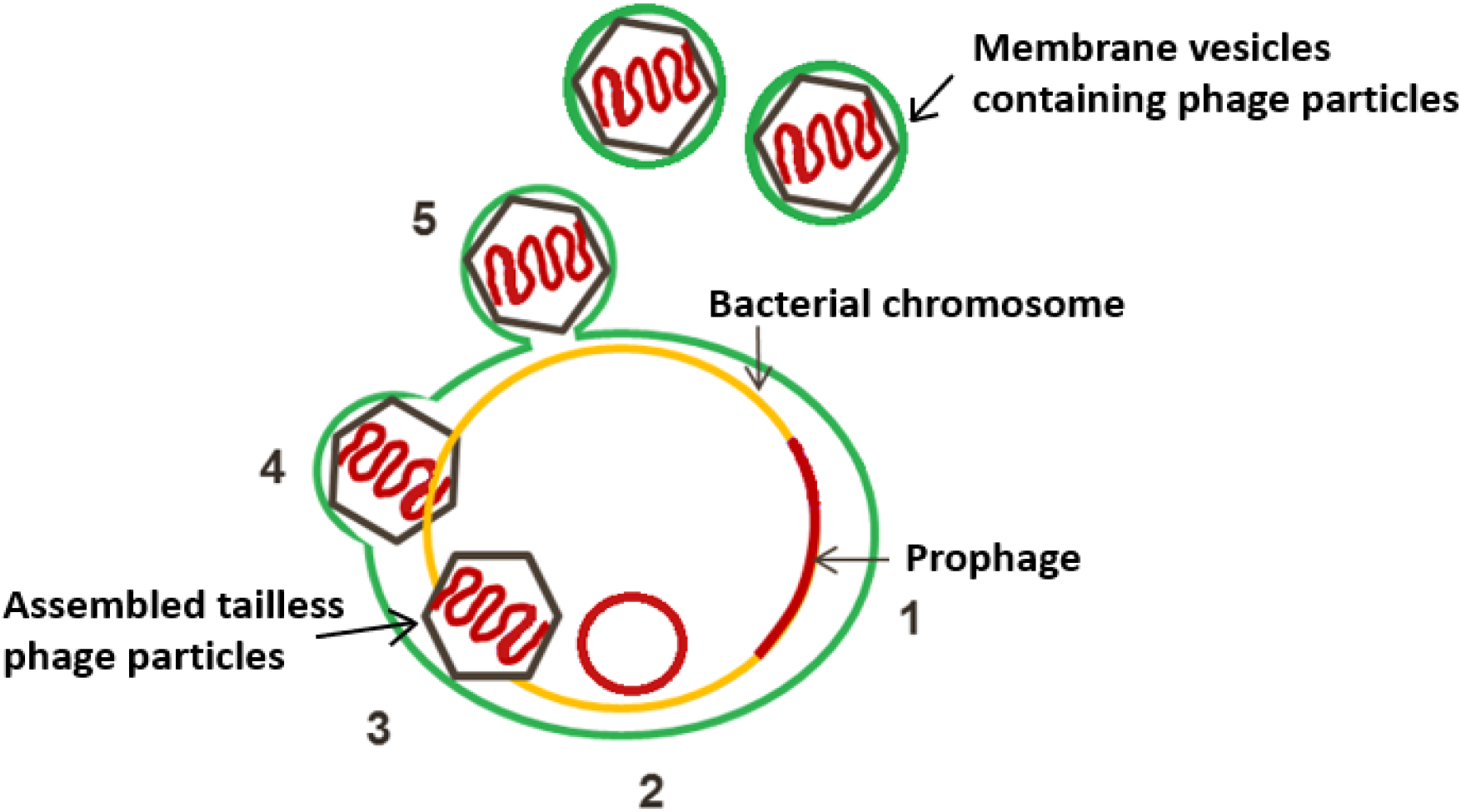
Schematic presentation of the proposed mechanism (step 1-5) of phage release from *Lactococcus lactis* TIFN1. Activation of proPhi1 (step 1 and 2) results in production of tailless *Siphoviridae* phage particles (3), enclosed in lipid membrane derived from the cytoplasmic membrane (green) (4), and released from the cells by a budding-like, non-lytic mechanism (5).

The prophage found in *L. lactis* TIFN1, referred to as proPhi1, is classified in the family of *Siphoviridae*, which members are by definition tailed bacteriophages. Genomic analysis also revealed that genes encoding tail structures are present in these prophages, but due to disruptions in some of the tail genes, the assembled phage particles show a tailless phenotype (6). Interestingly, the lipid-containing phages discovered so far, mostly assigned to families of *Corticoviridae*, *Cystoviridae*, *Plasmaviridae* and *Tectiviridae*, are exclusively tailless phages (42). Plausibly, the reason that the membrane-containing feature was not found in any tailed phages is that they already achieve successful infection with the help of the tail device that efficiently penetrates the cell envelop, and no alternative infection mechanism was required (42). Tailless phages, on the other hand, are evolved to utilize the membrane to infect or interact with their hosts. For example, enveloped phages use a membrane fusion mechanism to interact with the host and deliver their genetic materials (42–45). Therefore, it was part of the hypothesis that the tailless proPhi1 being enclosed in a lipid membrane could serve as an alternative infection strategy as the tail device is not available anymore, but so far we did not obtain evidence demonstrating the (re)infection of host by the membrane-enclosed tailless phage particles (data not shown). It remains to be investigated whether the hurdle was for membrane-enclosed phage particles to attach and enter the host, or rather for the tailless phage particles to inject their genetic material into the bacterial cytoplasm to complete the life cycle.

In previously described membrane-enclosed phages, the mechanism of incorporation of lipids to form virus-specific vesicles has been subjected to investigation. However, not everything is completely understood until today, but several mechanisms are proposed (42). For one, phage encoded membrane proteins trigger cytoplasmic membrane formation in the host, and enclose the phages during assembly in the cell. For example, *Cystoviridae* phage phi6 applies a mechanism, in which the protein P9 was found to facilitate cytoplasmic membrane formation in bacteria (46). Host-derived membrane components can be an alternative mechanism. In this case, phage-encoded membrane proteins are incorporated into the host membrane, providing a scaffold for phage assembly, and the assembled phage particles are released upon lysis of the host (47, 48). Examples are *Tectiviridae* phage PRD1 employing membrane protein P10 (49), and *Corticoviridae* phage PM2 employing membrane proteins P3 and P6 to interact with phage-specific areas on the cell membrane (42, 50). For all above mentioned phage-encoded proteins, we did not find homology to any of the proteins encoded on proPhi1 or any of the other prophages found in lactococci isolated from the starter culture Ur (6). However, it should be noted that other membrane-associated protein coding sequences were indeed predicted in the Ur prophages, namely ORF42 in proPhi1&5, ORF08 in proPhi2&4 and ORF49 in proPhi6 (6). Targeting phage particles to special areas of cell membrane by membrane-associated proteins could potentially be an explanation for the distinct lipid composition associated with released phage particles. Nevertheless, non-lytic release via a mechanism of budding is still not confirmed in other phages but suggested for plasmavirus (19) and is considered a very delicate life cycle of viruses, as it leaves the host alive while phages get to spread the progeny (42). Whether the non-lytic release of membrane-enclosed phages in *L. lactis* TIFN1 and other lactococcal strains found in the starter culture Ur is a result of long-term phage-host co-evolution thus becomes an even more interesting hypothesis, especially as we observed similar growth behavior during phage release in other Ur strains (23).

Another intriguing question is how the membrane-enclosed phage particles escape from the bacterial host without lysis, especially given the fact that *L. lactis* is Gram-positive, possessing thick cell wall outside the cell membrane. A similar question has been raised for extracellular membrane vesicles (MVs or EVs) produced by Gram-positive bacteria ever since the discovery of such phenomenon. It has already been known for a long time that Archaea, Gram-negative bacteria, and mammalian cells actively secrete the nano-sized, lipid bilayer-enclosed particles named EVs, harboring various nucleotide and protein cargos as a mechanism for cell-free intercellular interactions (51–53). Only recently, evidence was provided that EVs are also released by organisms with thick cell walls like Gram-positive bacteria, mycobacteria and fungi (54–57), but the mechanistic insights are still lacking. Brown et al. (2015) proposed several non-mutually exclusive mechanisms on the formation and release of EVs through thick cell walls, including the actions of turgor pressure, cell wall-modifying enzymes and protein channels. The most evidence-supported mechanism is via cell-wall modifying enzymes, namely autolysin (58) and prophage-encoded holin-endolysin (59, 60). Notably, phage particles have also been identified as part of the cargos in EVs produced by *Bacillus subtilis* (59). Further studies dedicated to elucidating the roles of autolysin and/or phage-encoded holin-endolysin in *L. lactis* TIFN1 would serve to reveal the release mechanism in this case.

Moreover, the effect of turgor pressure could also play a role in addition (57). It is plausible that upon prophage induction, the defective proPhi1 particles are abundantly assembled and accumulated in the cells, causing cytoplasmic crowding that results in elevated turgor pressure. The cell division site is often the target site of autolysins (61), in combination with induced phage-encoded endolysins, forming the weakest spot on the cell and giving opportunities for the phage particles to release under turgor pressure, which could explain our observation that the membrane-enclosed particles are mostly observed near the cell division sites, and have a distinct lipid profile comparing the whole cell samples. Therefore, we propose that the phenomenon of non-lytic membrane engulfed phage release observed in *L. lactis* TIFN1 could be driven by the concerted action of enzymatic activity and turgor pressure on the cell envelope, in combination with phage-encoded proteins to achieve phage-specific engulfment and release.

Although this is the first study to demonstrate non-lytic release of membrane-engulfed phages in LAB, we would like to point out that this could be a more common but up to now overlooked phenomenon in other microbial communities for two reasons. Firstly, studies focused on the detection of inducible prophages, use cell lysis/plaque formation as a benchmark for phage activation. Obviously, when (tailless) phage particles are released via membrane envelops or other non-lytic ways, no apparent phenotype will be observed thus discouraging further investigation. Secondly, it is a common practice in phage isolation protocol to employ chloroform to remove contaminating materials derived from bacterial cells (42), however, this treatment demolishes the membrane structures and therefore the lipid-containing phenotype is conceivably not retrieved in further analysis of phage particles. We hope that our findings will inspire further studies, not only in elucidating the detailed mechanism of this case, but also in awareness and discovery of similar phenomena in other microbial species, and further shedding light on bacteria-phage interaction and co-evolution.

## Data availability statement

The datasets used and/or analyzed during the current study are available from the corresponding author on reasonable request.

## Authors’ contributions

SA and EJS conceived the study. SA, EJS and YL designed the experiments. YL and SA executed the experiments and carried out the data analysis and interpretation. HB was involved in the chemical lipid analysis, JAGM prepared samples for electron microscopy and obtained the pictures. NY and SA did the fluorescent lipid detection. YL, SA, TA and EJS wrote the manuscript.

All authors read and approved the final manuscript.

## Acknowledgments

We thank Lieke Gijtenbeek and Jan Kok (RUG) for providing plasmids pCS1966, pVE6007, and pSEUDO-GFP. Emmanuelle Maguin (INRA, France) is acknowledged for providing pG^+^host9. The authors cordially thank Guido Staring (NIZO food research, Ede, the Netherlands) for performing LC-MS/ESI lipid analysis. The electron microscopy images were obtained with the help of Marcel Giesbers at the Wageningen Electron Microscopy Centre (WEMC) of Wageningen University (Wageningen, The Netherlands).

## Supplementary materials

**Figure S1.**
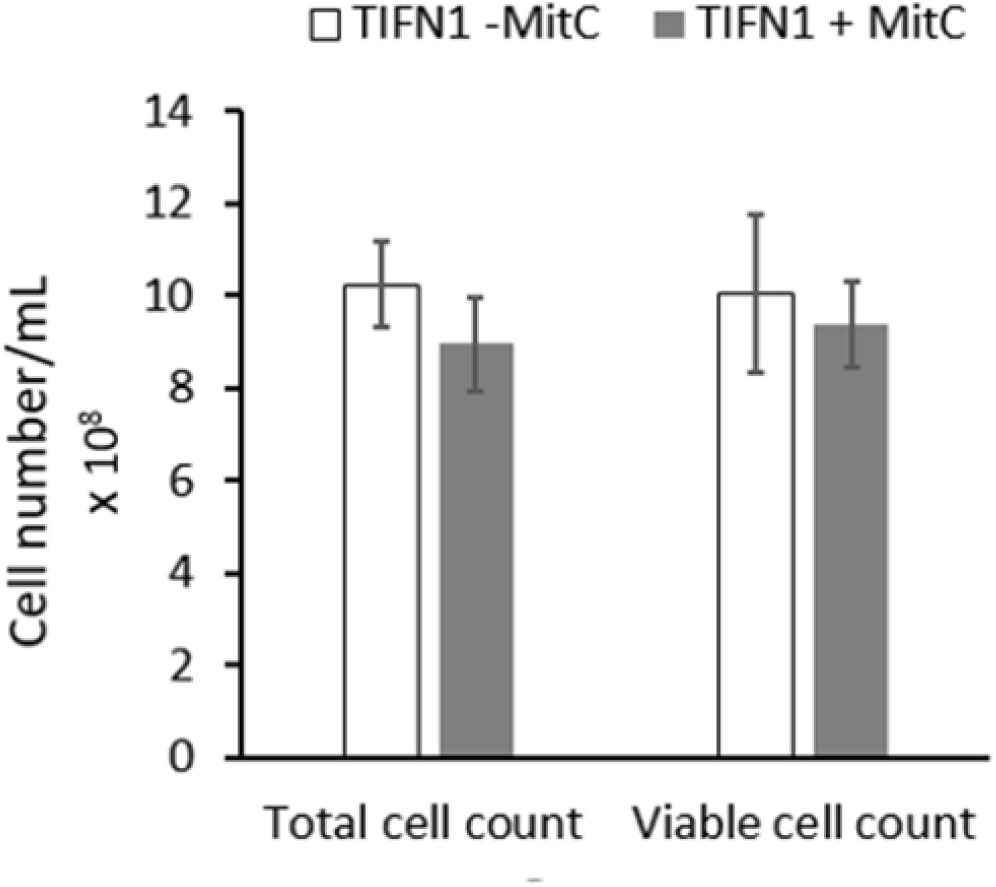
Cell count of *Lactococcus lactis* strain TIFN1 under phage induction conditions. Total cell count (obtained by counting cells using a haemocytometer) and viable cell count (determined by plating and colony count) in TIFN1 cultures induced with MitC at 7 hours and control cultures without induction.

**Figure S2.**
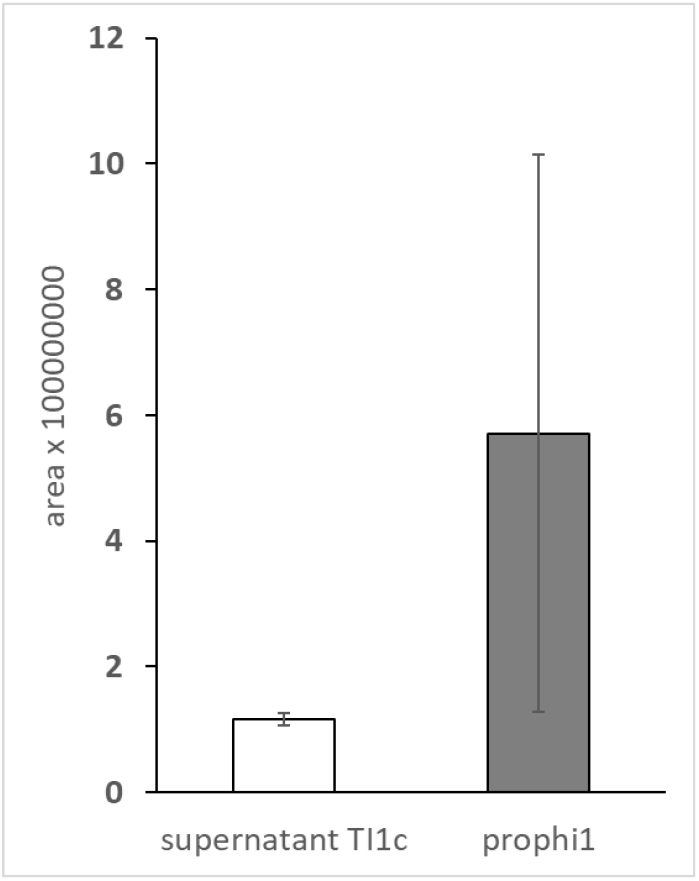
Lipid (sum of phosphatidyl glycerol and cardiolipin) signal detected in culture supernatant of phage-free control TI1c and proΦ1 collected from culture supernatant of TIFN1.

